## Supplemental Materials for "Compartmentalization and persistence of dominant (regulatory) T cell clones indicates antigen skewing in juvenile idiopathic arthritis"

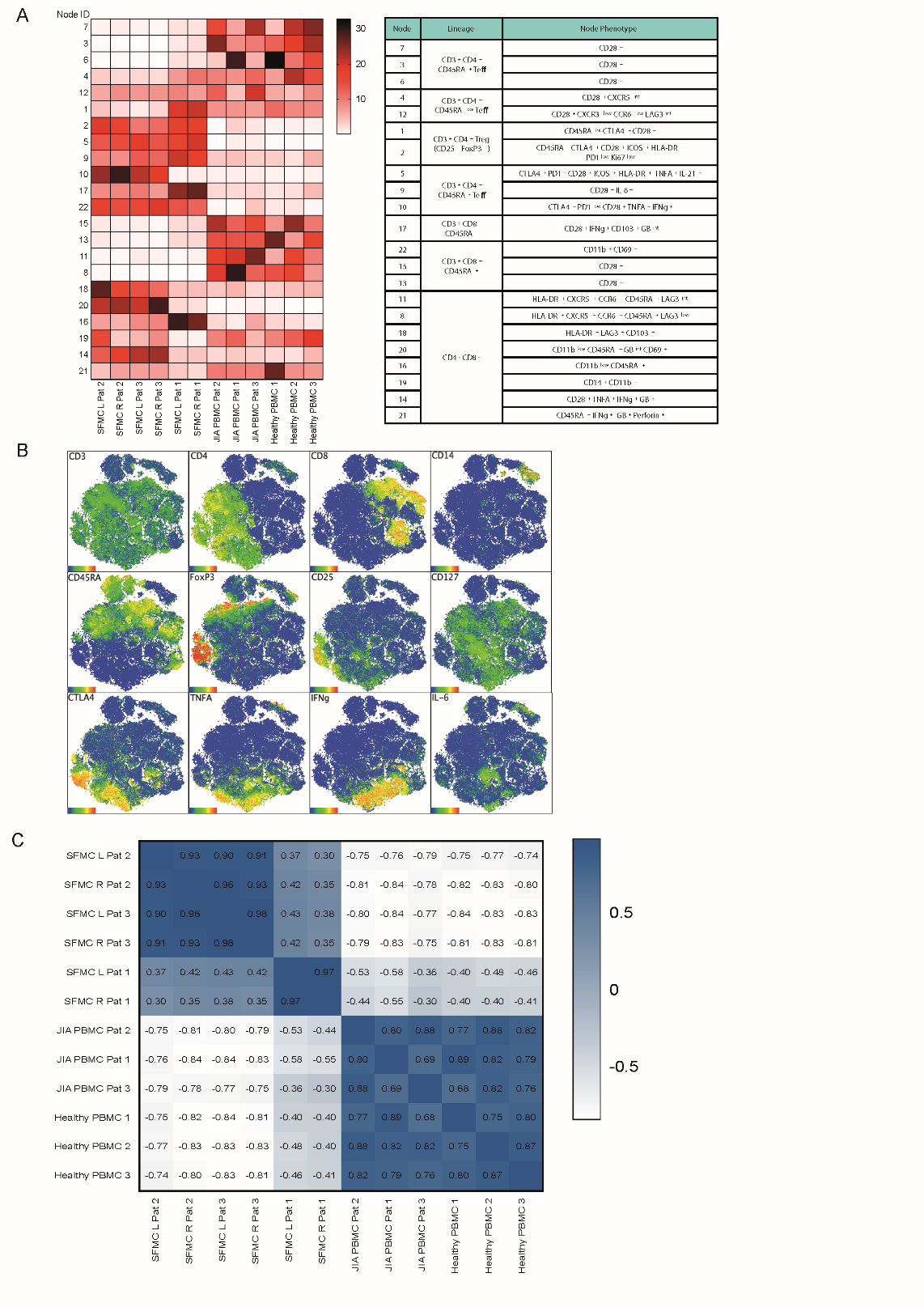

**Supplemental Figure 1.** **Preliminary analysis reveals correlation between SFMC from distinct joints. A** Node frequency showing the distribution of T cell markers across the nodes of SFMCs an PBMCs in the CyTOF analysis. **B** Marker expression of t-SNE dimensional reduction and k-means clustering analysis on SFMC and PBMC samples **C** Correlation matrix using spearman correlation of the entire spectrum of node frequency given in A.

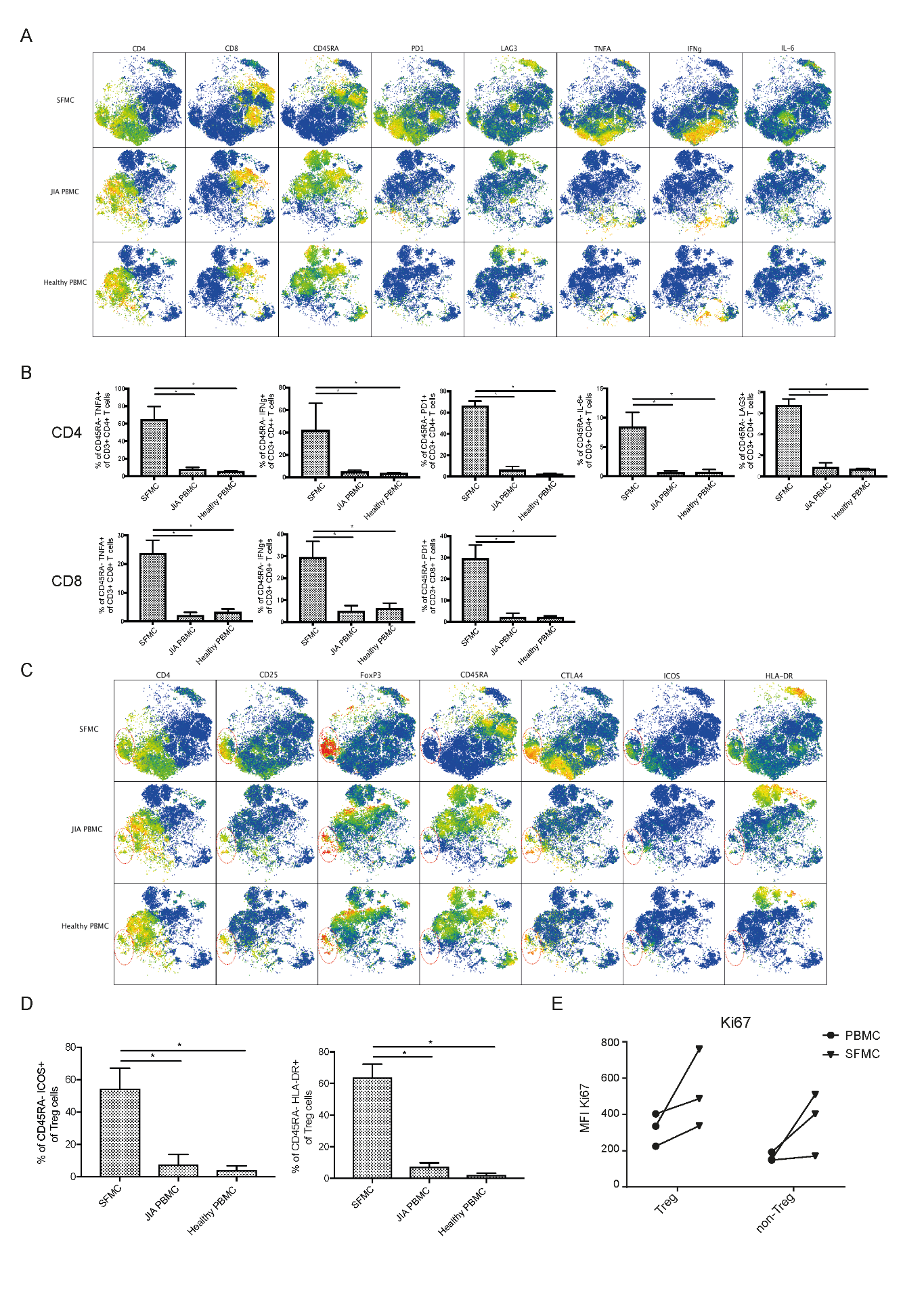

**Supplemental Figure 2. SFMC display an activated expression profile. A** T-SNE plots showing the expression profile of phenotypical and functional markers in SFMC, PBMC from JIA patients and PBMC from healthy children. **B** Bar charts showing the percentage of specific cell populations within CD4+CD45RA- and CD8+CD45RA- cells (non-parametric Mann-Whitney, * = p <0.05). **C** T-SNE plots showing the expression profile of phenotypical and functional Treg markers in SFMC, PBMC from JIA patients and PBMC from healthy children. **D** Quantification of CD45RA-ICOS+ and CD45RA-HLA-DR+ expression on CD25+ FOXP3+ Treg (non-parametric Mann-Whitney, * = p <0.05). E MFI of Ki67 protein expression in Treg and non-Treg as determined by flow cytometry**.**

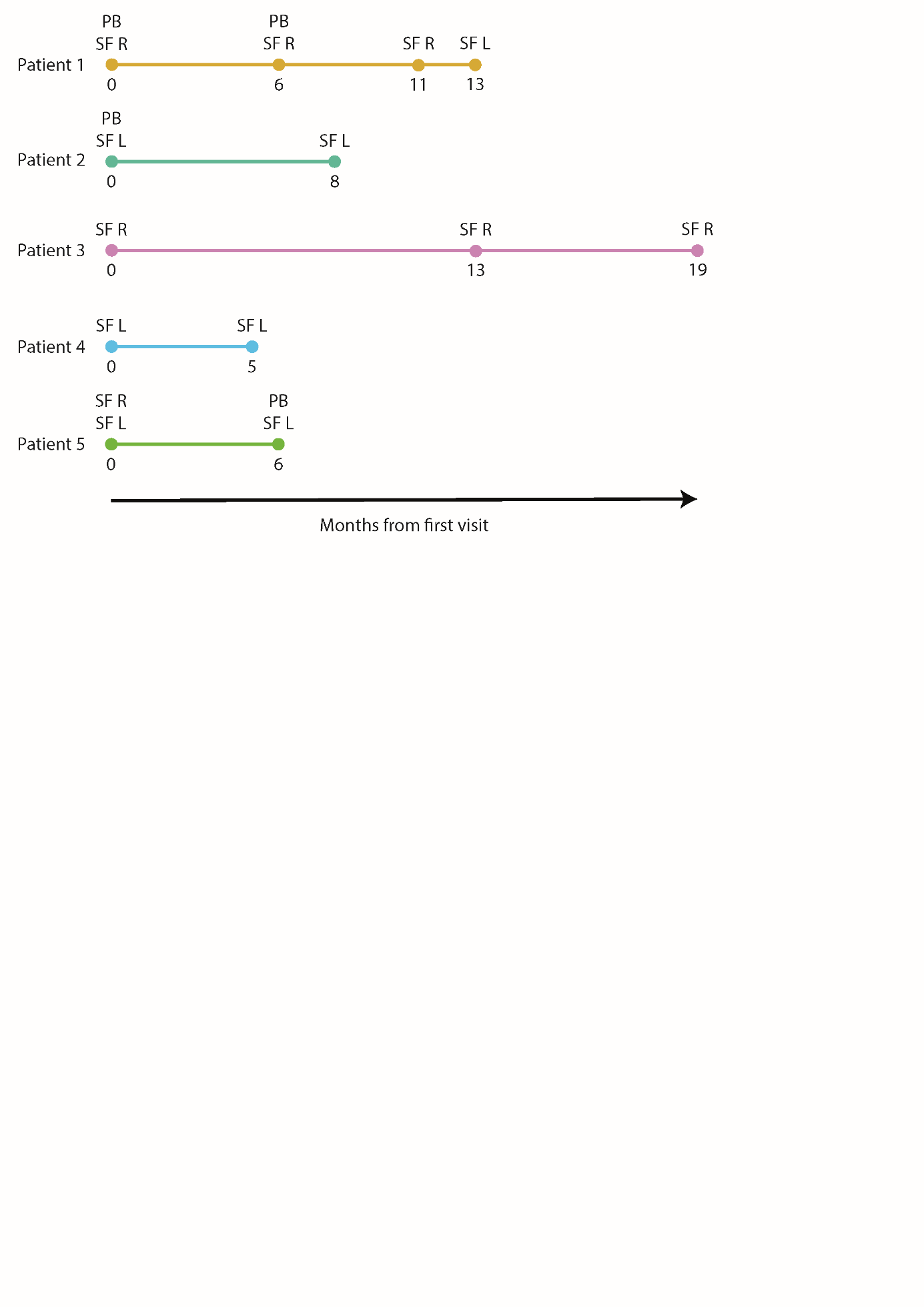

**Supplemental Figure 3. Longitudinal sampling timelines of JIA patients.** PB = peripheral blood, SF = synovial fluid, L = left, R = right.

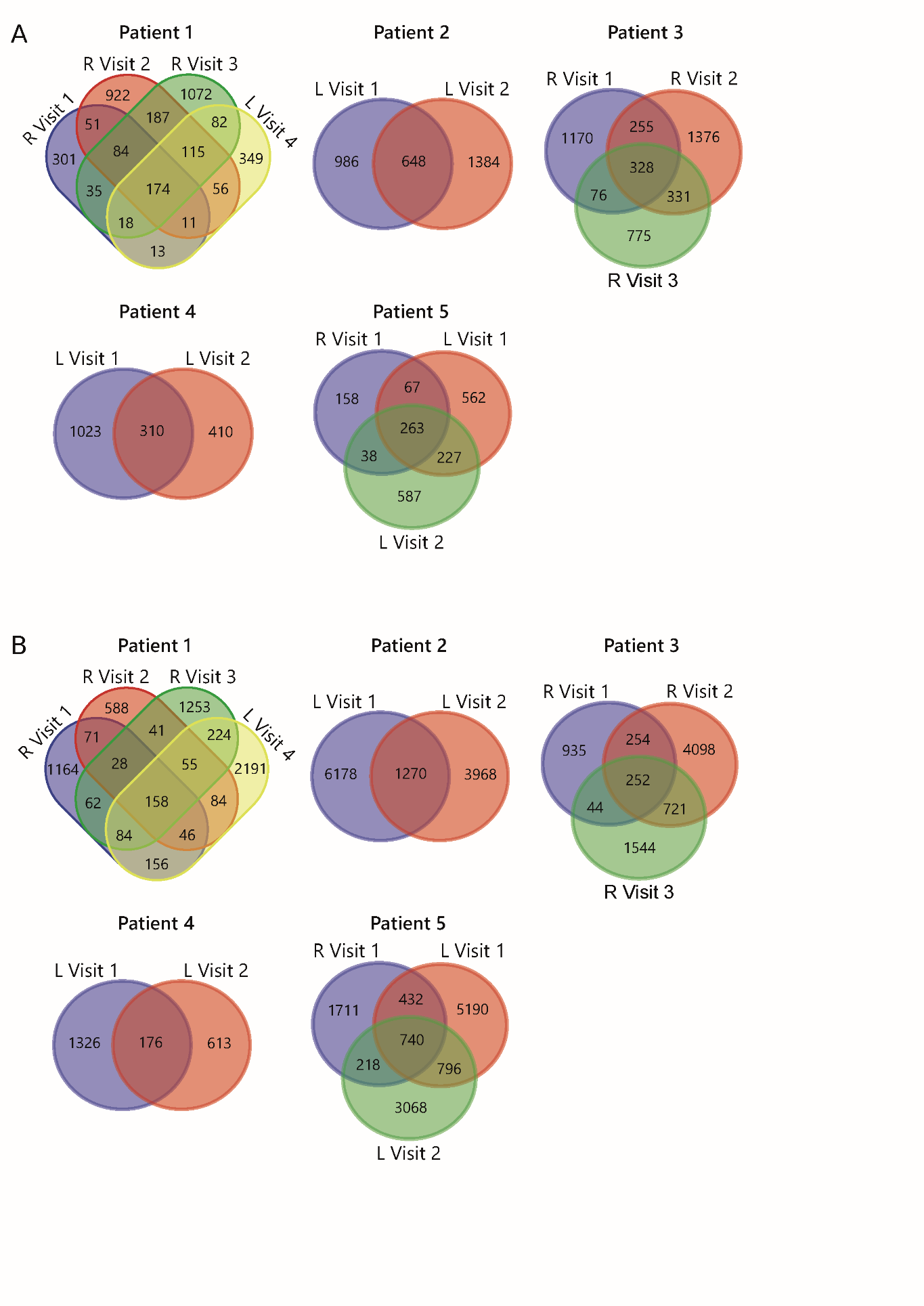
**Supplemental Figure 4. TCR overlap analysis. A** Venn diagrams displaying the overlap of all unique TCRβ clones, defined by amino acid sequence, for longitudinal SF samples from all patients for Tregs and **B** non-Tregs.

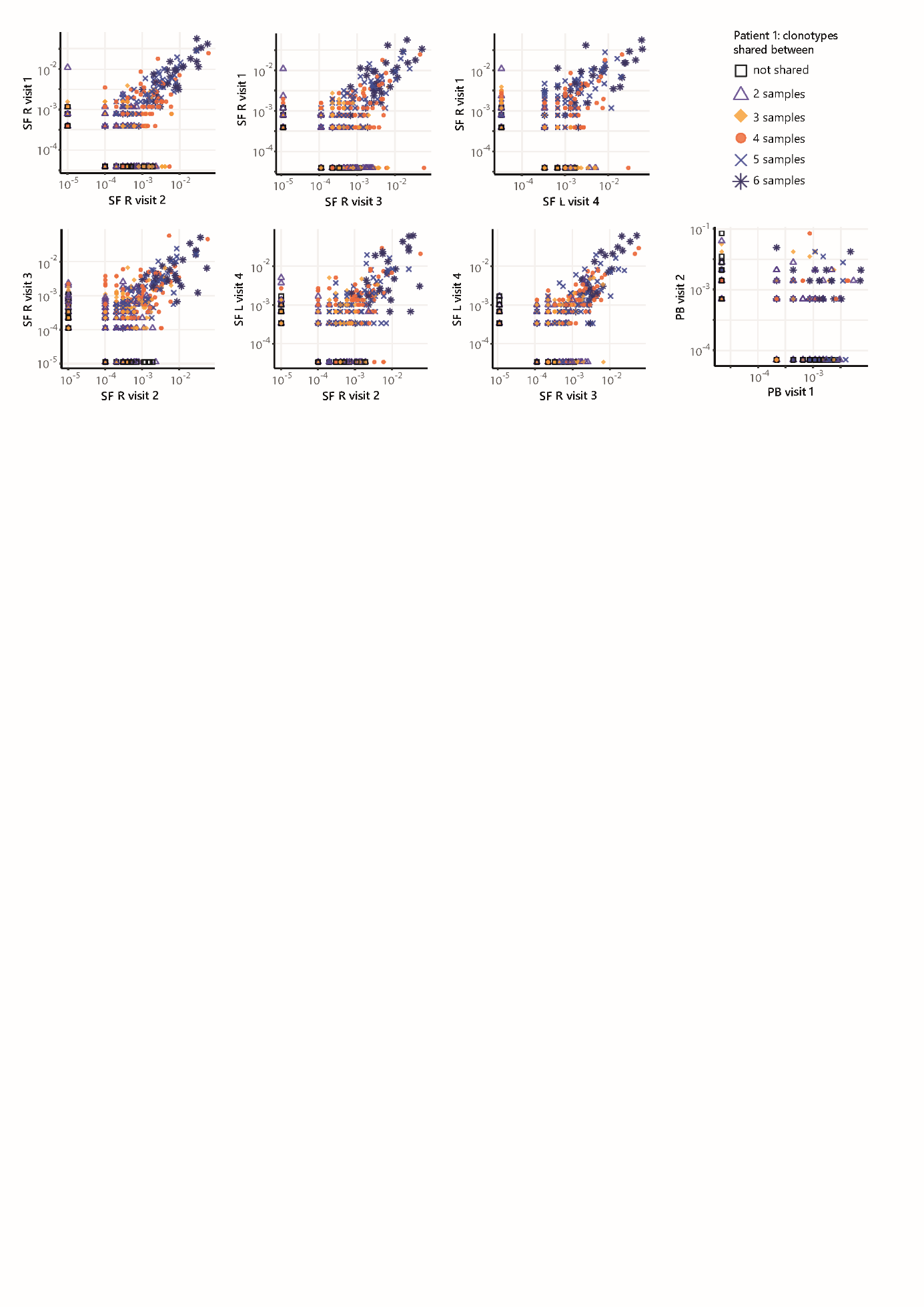

**Supplemental Figure 5. Frequencies of TCRs from persistent Tregs shared across SF and PB samples.** Frequency plots showing the overlapping Treg clones between visits for SF and PB, with color coding and shapes highlighting the number of samples in which unique clones are found.

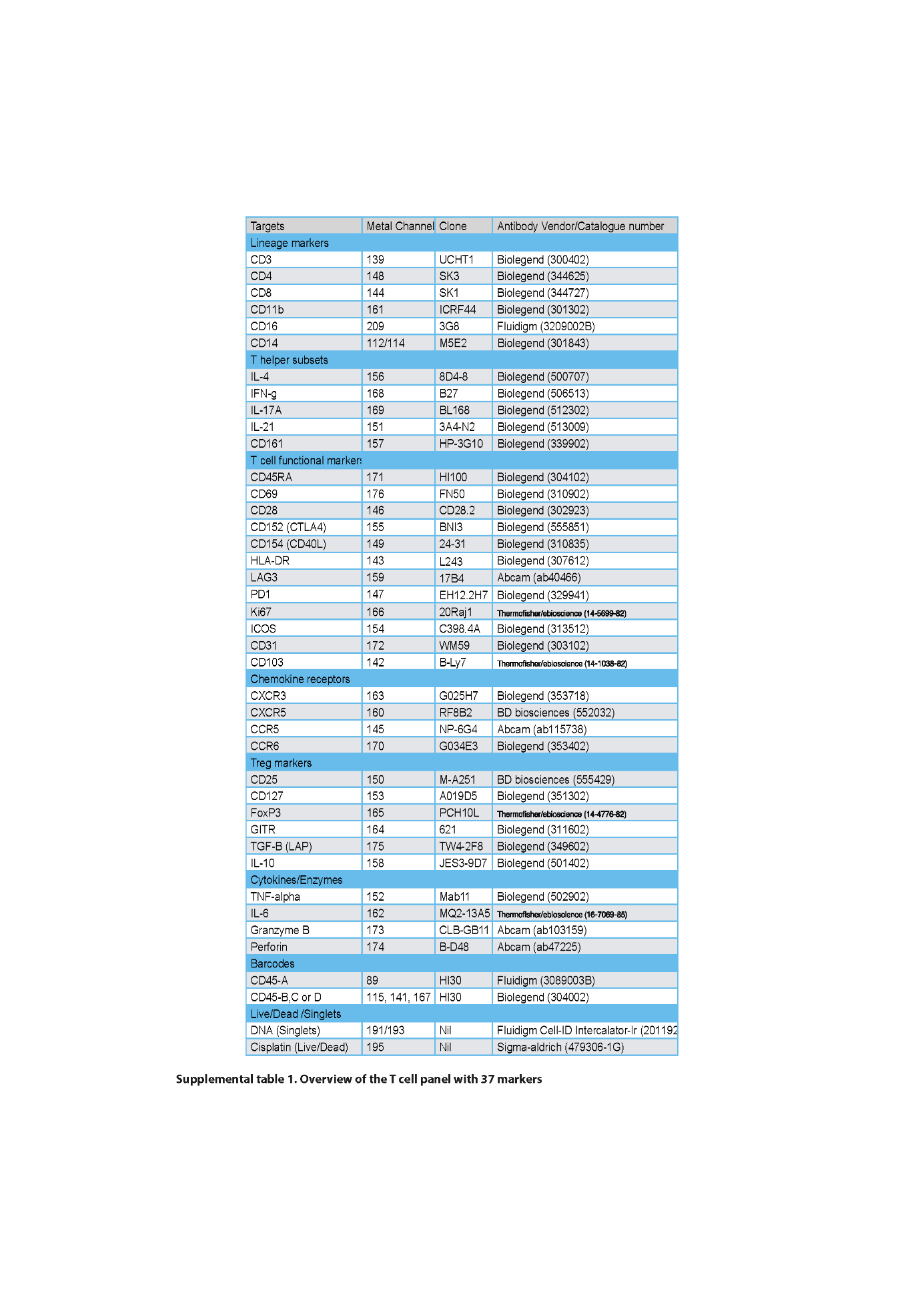

**Supplemental Table 1. Overview of the T cell panel used for CyTOF with 37 markers.**
